# G-OnRamp: Generating genome browsers to facilitate undergraduate-driven collaborative genome annotation

**DOI:** 10.1101/781658

**Authors:** Luke Sargent, Yating Liu, Wilson Leung, Nathan T. Mortimer, David Lopatto, Jeremy Goecks, Sarah C. R. Elgin

**Affiliations:** Department of Biomedical Engineering, Oregon Health & Science University, Portland, Oregon, United States of America; Department of Biology, Washington University in St. Louis, St. Louis, Missouri, United States of America; School of Biological Sciences, Illinois State University, Normal, Illinois, United States of America; Department of Psychology, Grinnell College, Grinnell, Iowa, United States of America

## Abstract

Scientists are sequencing new genomes at an increasing rate with the goal of associating genome contents with phenotypic traits. After a new genome is sequenced and assembled, structural gene annotation is often the first step in analysis. Despite advances in computational gene prediction algorithms, most eukaryotic genomes still benefit from manual gene annotation. Undergraduates can become skilled annotators, and in the process learn both about genes/genomes and about how to utilize large datasets. Data visualizations provided by a genome browser are essential for manual gene annotation, enabling annotators to quickly evaluate multiple lines of evidence (*e.g.*, sequence similarity, RNA-Seq, gene predictions, repeats). However, creating genome browsers requires extensive computational skills; lack of the expertise required remains a major barrier for many biomedical researchers and educators.

To address these challenges, the Genomics Education Partnership (GEP; https://gep.wustl.edu/) has partnered with the Galaxy Project (https://galaxyproject.org) to develop G-OnRamp (http://g-onramp.org), a web-based platform for creating UCSC Assembly Hubs and JBrowse genome browsers. G-OnRamp can also convert a JBrowse instance into an Apollo instance for collaborative genome annotations in research and educational settings. G-OnRamp enables researchers to easily visualize their experimental results, educators to create Course-based Undergraduate Research Experiences (CUREs) centered on genome annotation, and students to participate in genomics research.

Development of G-OnRamp was guided by extensive user feedback from in-person workshops. Sixty-five researchers and educators from over 40 institutions participated in these workshops, which produced over 20 genome browsers now available for research and education. For example, genome browsers for four parasitoid wasp species were used in a CURE engaging 142 students taught by 13 faculty members — producing a total of 192 gene models. G-OnRamp can be deployed on a personal computer or on cloud computing platforms, and the genome browsers produced can be transferred to the CyVerse Data Store for long-term access.

## Introduction

### The need for G-OnRamp

A considerable effort has been made over the last two decades to improve undergraduate science education by engaging students in the process of science, as well as acquainting them with the resulting knowledge base. For the life sciences these efforts were perhaps best enunciated by the AAAS report *Vision and Change in Undergraduate Biology Education* [1]. One of the strategies found to be effective in engaging large numbers of undergraduates in doing science is the CURE, or Course-based Undergraduate Research Experience ([2]; see [3] and [4] for examples). Within computational biology, a number of groups have found that genome annotation is a research problem that can be adapted to this purpose.

With the decreasing cost and wide availability of genome sequencing [5], the bottleneck for utilizing genomics datasets to address scientific questions is shifting from the ability to produce data to the ability to analyze and interpret data. Genome annotation—labeling functional regions of the genome such as gene boundaries, exons, and introns—benefits from a combination of computational and manual curation of data. With appropriate tools and training, undergraduates can make a significant contribution to a community annotation project, where scientists work together to annotate an entire genome. Gene annotation builds on what students are learning about gene structure, while requiring them to grapple with multiple lines of evidence to establish defendable gene models. Student annotation projects thus are mutually beneficial for researchers and for students, enabling unique science and providing a multi-faceted learning experience for students [6, 7, 8, 9, 10].

However, despite the improvements in tool accessibility and quality, there remain technical barriers that must be overcome to perform genome annotation. Many biology researchers and educators lack detailed knowledge of informatics and computational tools. When these scientists acquire the genome assembly of their favorite organism, a major barrier is the need to use multiple bioinformatics tools to analyze the genome assembly and visualize the results in a genome browser — the display tool central to community annotation. There are several good options, but most either require substantial computer skills and bioinformatics expertise to use, or have compute and storage limits that restrict the size/complexity of genome assemblies that can be analyzed using the platform [11, 12, 13, 14, 15].

We developed G-OnRamp to address these concerns. G-OnRamp is a collaboration between the Galaxy project (https://galaxyproject.org/), an open-source, web-based computational workbench for analyzing large biological datasets [16], and the Genomics Education Partnership (GEP; http://gep.wustl.edu/) [8, 17]. Among G-OnRamp’s principal goals is lowering technical barriers to enable biologists to construct either a UCSC Assembly Hub [18] or a JBrowse/Apollo genome browser [19]. G-OnRamp accomplishes this by providing a collection of tools, workflows and services pre-configured and ready to process data and enable annotation. Students, educators and researchers can bypass most of the system administration tasks involved in generating a genome browser and focus on using the genome browser to address scientific questions. Our assessment results in the classroom demonstrate that the genome browsers produced by G-OnRamp are effective tools for engaging undergraduates in research and in enabling their contributions to the scientific literature in genomics.

## Results

### Overview of the components

#### Genome annotation needs for the Genomics Education Partnership

The GEP is a consortium of faculty members from over 100 educational institutions, which annually introduces more than 1300 undergraduates to genomics research through engagement in collaborative annotation projects (Fig 1A). The GEP core organization provides technical infrastructure as well as identifying research questions that would benefit from high quality gene annotations, particularly those where utilizing comparisons across multiple species can provide insights. By engaging the talents of “massively parallel undergraduates,” one can gather data (high quality annotations of hundreds of genes) that could not be obtained otherwise, given the high labor costs. To ensure that the gene annotations are high quality, each gene is annotated by at least two students working independently, and the results are reconciled by experienced students (Fig 1B).

**Fig 1.**
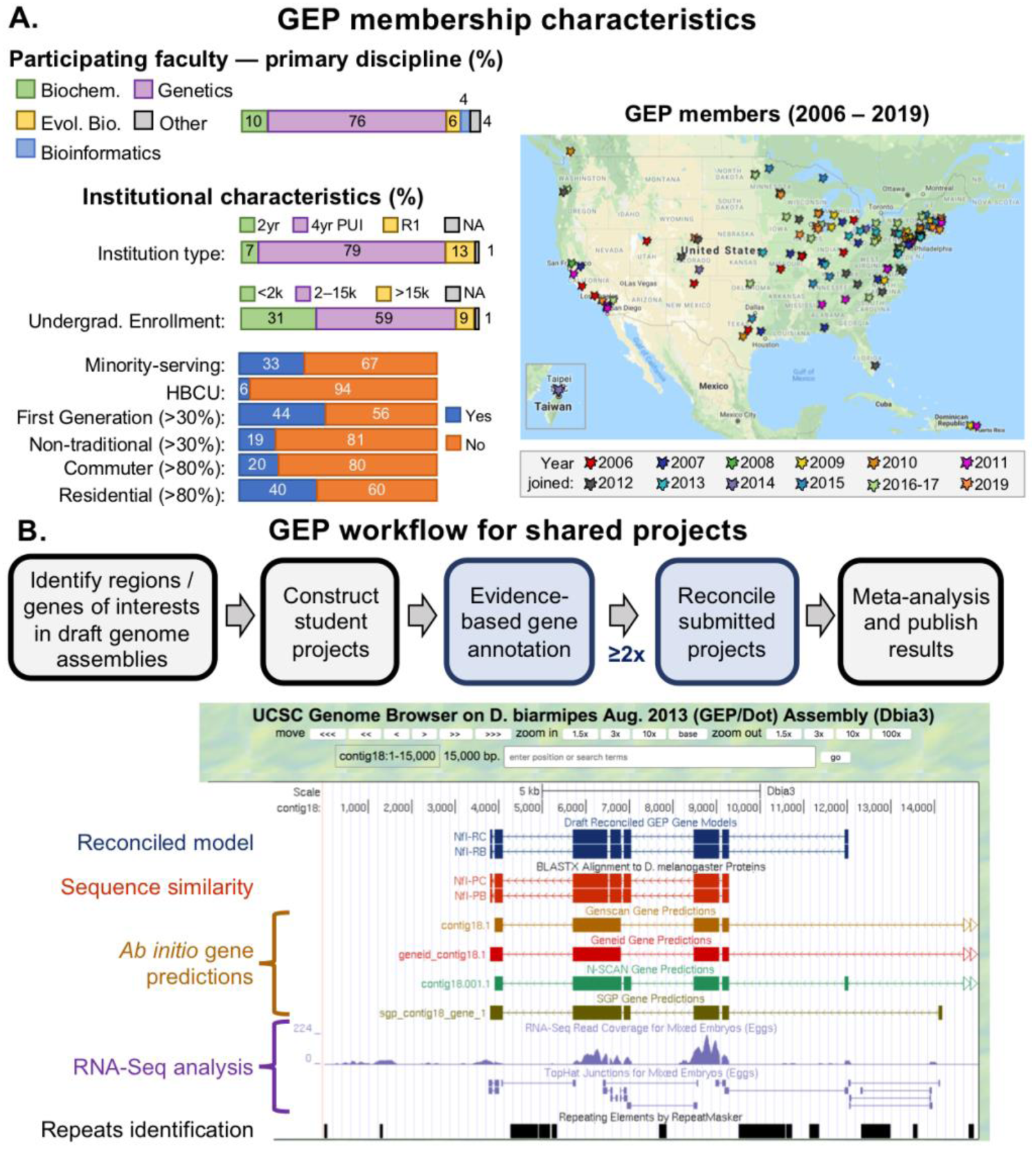
Overview of the Genomics Education Partnership. A. Membership characteristics: participating faculty primarily teach genetics (although other disciplines are represented), and most often teach at Primarily Undergraduate Institutions (PUIs) across the USA; faculty at community colleges and R1 research universities also participate. The geographical distribution of member schools and year of joining GEP are shown on the map. The member schools serve a diverse undergraduate student body, with 33% Minority-Serving Schools, including six HBCUs (Historically Black Colleges and Universities); 44% of the schools have 30% or more first-generation students, 11% have 30% or more non-traditional students (over 25 yrs of age), and 20% are commuter schools, with over 80% of the students commuting. See the Current GEP Members page (http://gep.wustl.edu/community/current_members) for a complete list of participating faculty with their schools. B. Students in the GEP work together to produce high-quality annotation of a genome region or a collection of genes of interest identified by a Lead Scientist. “Student projects” are provided as genome browser pages (see lower portion of the figure) with from one to seven potential genes (and other features of interest) for annotation. Browser tracks show available evidence for a gene, including gene conservation (Sequence similarity track and additional BLAST searches), presence of large open reading frames and other appropriate signals (*ab initio* gene predictions), and evidence of gene expression (RNA-seq data, Top-Hat analysis results, etc.). Students work from these multiple lines of evidence, some of which may initially appear contradictory, to generate a gene model that they can defend. In the case shown, the sequence similarity search (BLAST) failed to identify putative upstream exons, whose presence is supported by RNA-seq data and Top-Hat analysis. Students take responsibility for the workflow steps shown in light blue, while the Lead Scientist’s research group is responsible for the steps shown in grey. Pre-/post course assessment has shown the effectiveness of such a collaborative annotation project both for supporting student learning about genes and genomes and in providing a research experience [17, 20, 21].

These collaborative genome annotation projects can be performed by students using either a genome browser or a genome annotation editor such as Apollo. Pedagogically, there are advantages to requiring students to initially examine the evidence tracks on a genome browser, using the data to determine the precise exon coordinates for their gene model, and recording the results in an Excel worksheet or other table. These models can then be imported into the genome browser as custom tracks, and used as evidence in the final reconciliation. Currently, the GEP uses a hybrid approach, whereby students in GEP courses use a UCSC Genome Browser to construct the initial gene models, while experienced students use the Apollo annotation editor for finale reconciliation. See Fig 2 for an example of a typical error in a gene model submitted by a GEP student, viewed in Apollo for reconciliation. Overall, we see complete agreement in 60% – 80% of the gene models submitted, depending on the difficulty of the project.

**Fig 2.**
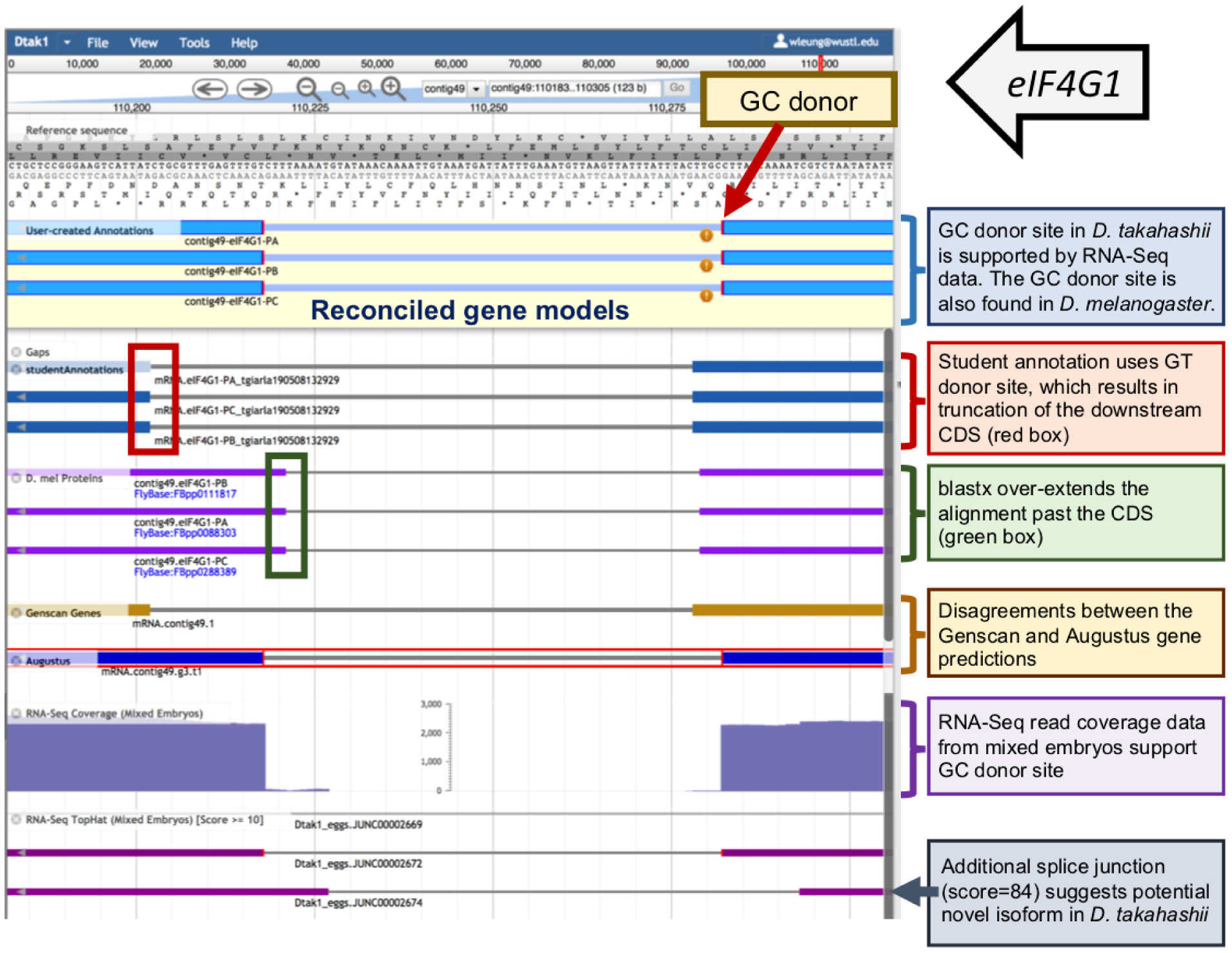
Apollo overview. After uploading data to Apollo via G-OnRamp’s “Create or Update Organism” tool, a user can choose which tracks to display with computational and experimental evidence, including submitted annotations from students, and begin to create her own gene model in a user-created annotations panel. Pictured is the Apollo interface showing provided sample data and computed lines of evidence, in addition to student annotation data and the final reconciled gene models (shown in the annotations panel). The genome browser image illustrates a typical error by one student annotator at an intron/exon boundary, and the reconciled model generated by an experienced student annotator. Based on RNA-Seq data and the use of the non-canonical GC donor site in the informant species (*Drosophila melanogaster*), the reconciled gene model for the *D. takahashii* ortholog of *eIF4G1* uses a non-canonical GC splice donor site instead of the GT donor site proposed by the student annotator.

GEP faculty have worked collaboratively to generate and maintain curricula to introduce students to the appropriate computer-based tools and to the scientific questions under study [8, 20]; all such materials are available on the GEP website under a “creative commons” license. Students who contribute documented gene models, and participate in reading and critiquing the final manuscript, are co-authors on the resulting scientific publication (*e.g*., [22], [23]). G-OnRamp was conceived by the GEP as a component of the technical infrastructure, simplifying the process of generating genome browsers. This capability should allow biology faculty to diversify the research questions under study, exploiting newly sequenced genomes as they become available.

#### G-OnRamp overview

G-OnRamp is a Galaxy-based analysis platform providing a collection of tools and services that enable collaborative genome annotation in an efficient, user-friendly, and web-based environment (http://www.g-onramp.org; [24]). Galaxy is used across the world by thousands of scientists, and one of its key features is a web-based user interface that anyone can use for complex biological analyses regardless of their computational knowledge. G-OnRamp is configured with tools for sequence similarity searches, gene predictions, RNA-Seq data analysis, and repeat analysis (Fig 3). These tools are combined into multi-step workflows that process a target genome assembly and create a UCSC Assembly Hub (which can be viewed at the official UCSC Genome Browser; http://genome.ucsc.edu) or a locally-bundled JBrowse instance. G-OnRamp also provides tools to import a JBrowse instance into Apollo to facilitate real-time collaborative genome annotation (https://genomearchitect.readthedocs.io/en/latest/; [10]). In a pedagogical example, an instructor can deploy G-OnRamp, upload the data, run a workflow to generate a JBrowse genome browser for visualization, and use the G-OnRamp Apollo interaction tools to convert the genome browser hub to Apollo for collaborative analysis by students.

**Fig 3.**
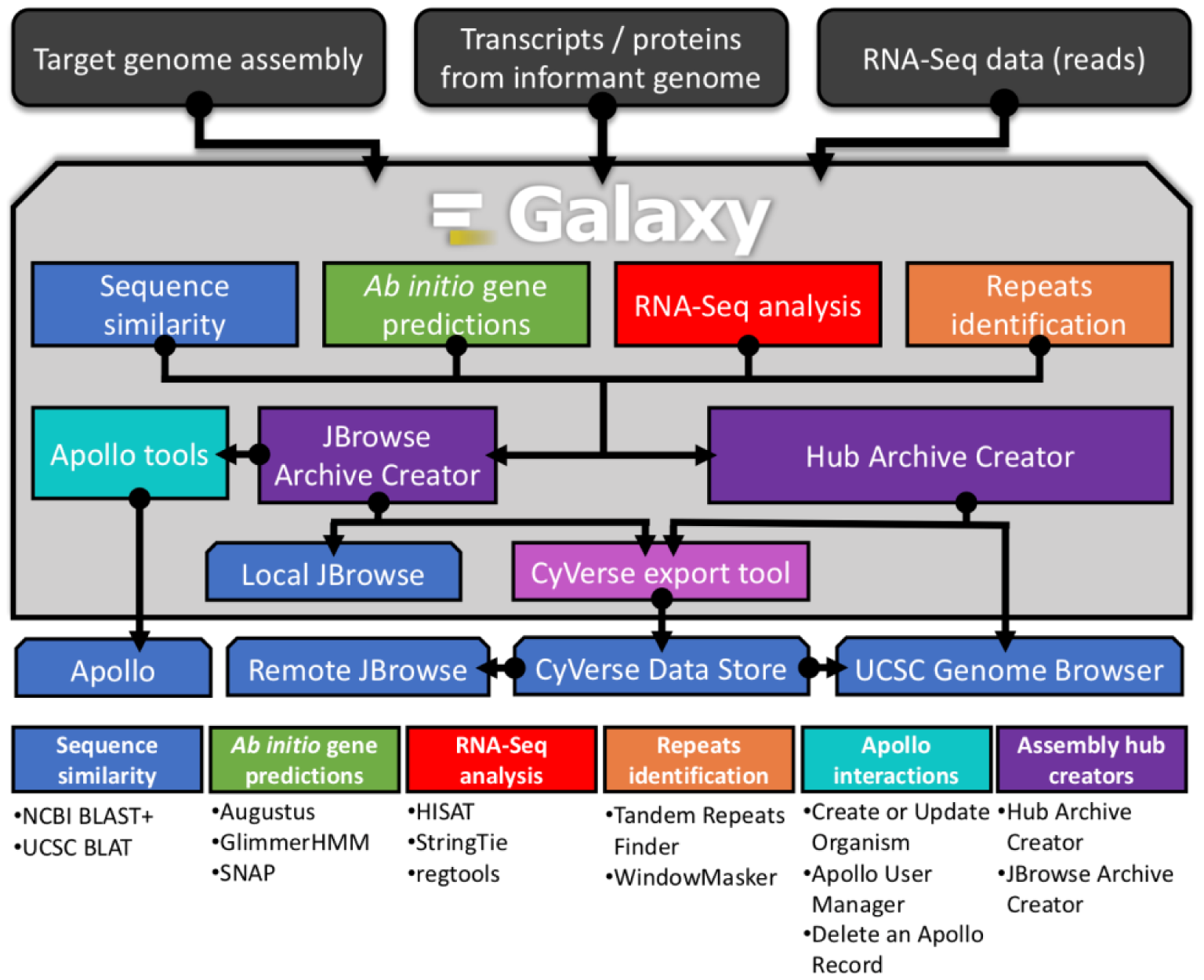
G-OnRamp overview. G-OnRamp is a Galaxy-based platform with analysis workflows that process a target genome assembly, transcripts and proteins from an informant genome, and RNA-Seq data from the target genome to create a genome browser for individual or collaborative annotation. Four sub-workflows (sequence similarity, *ab initio* gene predictions, RNA-Seq analysis, and repeats identification) run concurrently and generate the data for manual gene annotation. Data produced by the sub-workflows is used by the Hub Archive Creator (HAC) tool to create UCSC Assembly Hubs and by the JBrowse Archive Creator to create JBrowse genome browsers. The Apollo interaction tools convert JBrowse genome browsers into an Apollo instance to facilitate collaborative annotations. Genome browsers produced by G-OnRamp can be transferred to the CyVerse Data Store via the CyVerse export tool for long-term storage and visualization. The “Tool Suites” panel (below) lists the primary tools in each sub-workflow and the tools provided by G-OnRamp to create and manage Apollo instances. See [24] and http://g-onramp.org for further details.

#### Overview of genome annotation with Apollo: efficiency and crowd management

Apollo was included in G-OnRamp as it substantially increases the efficiency of gene annotation. Using Apollo, students can dynamically interact with evidence tracks, selecting the desired exons (by drag and drop) for assembly into a gene model. With effective permission management, annotation can be done separately (different students annotating different genes), iteratively (annotated genes being passed from one student to another) or simultaneously (students collaborate to annotate the same gene at the same time).

To aid permission-driven access control, G-OnRamp provides interaction tools (based on tools developed by the Galaxy community; [25]) for managing user accounts and genome assemblies in an Apollo instance. For example, a G-OnRamp administrator can use the “Create or Update Organism” tool to create a new Apollo instance or modify an existing Apollo instance. The Apollo User Manager tool provides fine-grained access controls; an administrator can control the read, write, and export permissions of individual users or groups of users. For example, instructors can use the Apollo User Manager to create accounts for a group of students enrolled in a course, and to limit their access to a subset of the genome assemblies in the Apollo instance.

### Using G-OnRamp in research and education settings

#### G-OnRamp workshops and evaluation

To grow the community of users and better tailor G-OnRamp to their needs, we hosted two beta-testers workshops in 2017 and two “train the trainer” workshops in 2018 to introduce researchers and educators to the platform. The goal of these workshops was to familiarize members of the community with G-OnRamp and to solicit feedback. These workshops attracted 53 diverse participants from over 40 institutions across the world, demonstrating that G-OnRamp satisfies a need for both researchers and educators alike (Fig 4).

**Fig 4.**
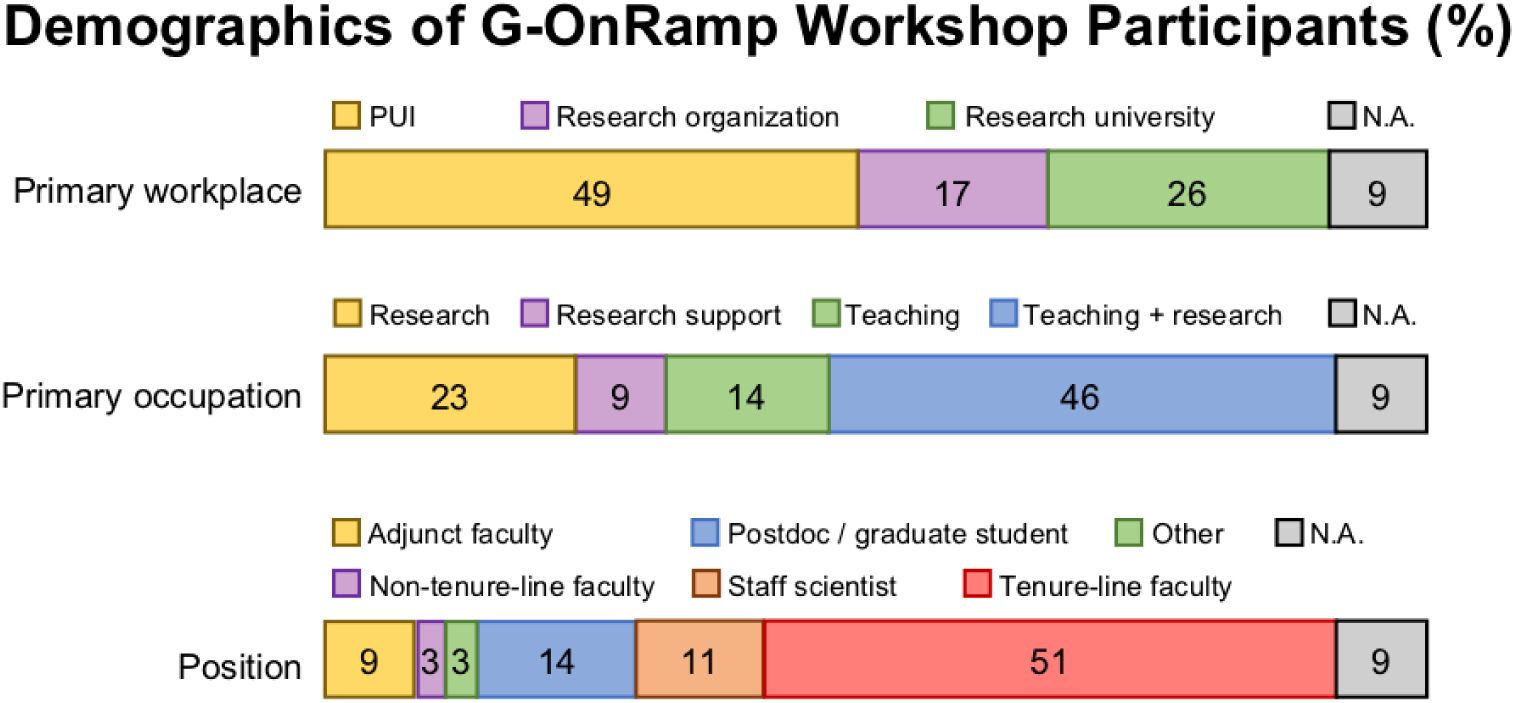
Demographics of G-OnRamp workshop participants. Of the 53 workshop participants eligible, 35 responded to the demographics questions (response rate = 66.0%). Many G-OnRamp workshop participants are tenure-line faculty members who work at primarily undergraduate institutions (PUIs), where they are involved in both teaching and research. Other participants focus mainly on research, either carrying out research or providing research support.

In addition to following a general training curriculum (available at http://g-onramp.org/training) on sample data, attendees were encouraged to bring their own genome assembly for processing and genome browser hub creation. Over 20 publicly-available genome browsers were created by workshop participants and the users that tested prototype G-OnRamp versions. Browsers generated during the 2017 and 2018 workshops demonstrate results obtained for genomes with assembly sizes ranging from 70Mb to 2.1Gb and with scaffold counts ranging from 53 to 271,888 (Table 1A). These genome browsers are hosted on the CyVerse Data Store [26] and are available via the “View Genome Browser” button on the G-OnRamp website (http://g-onramp.org/genome-browsers).

**Table 1A.**
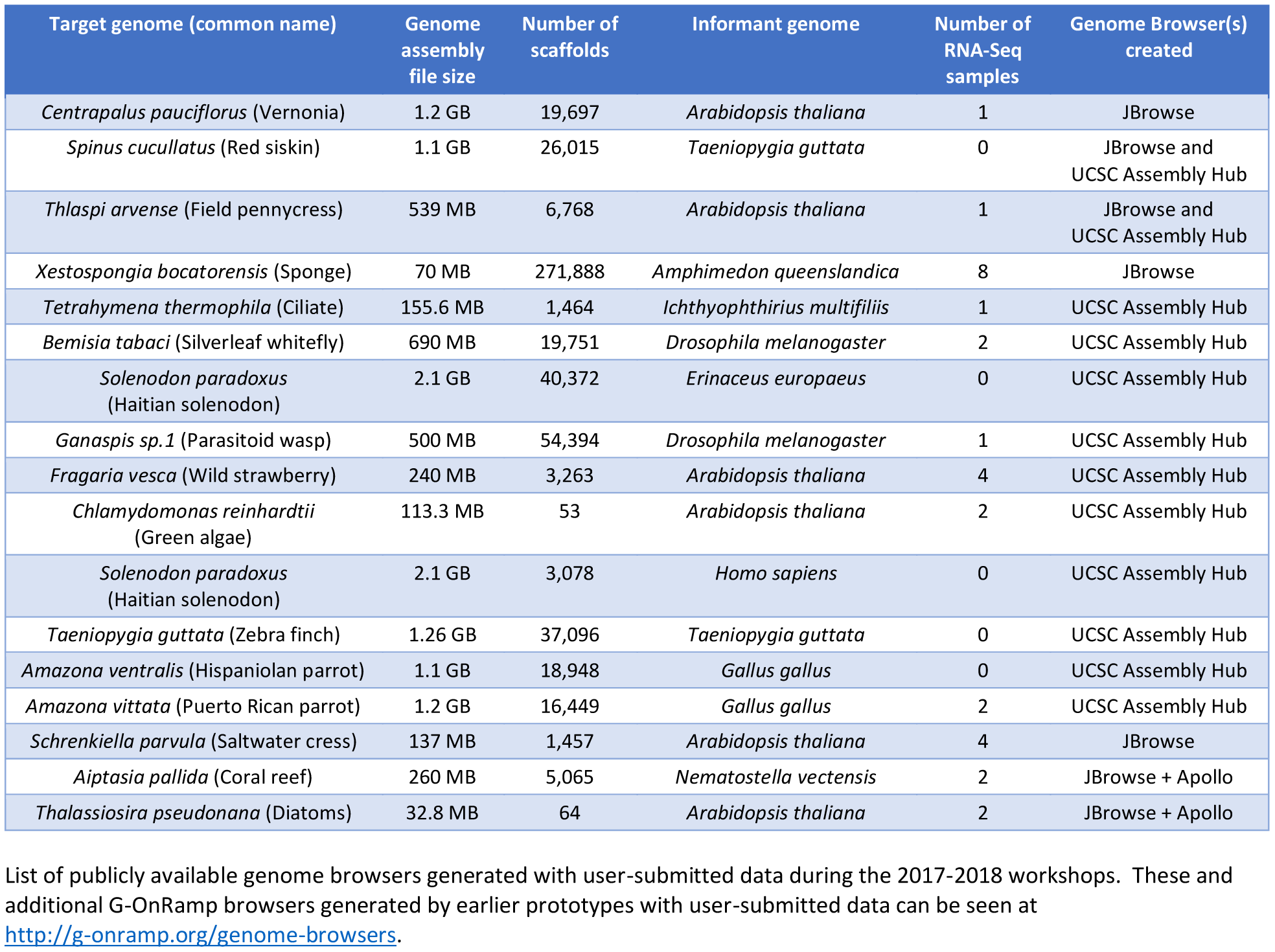
Publicly available genome browsers.

#### G-OnRamp features

Feedback collected from participants after each workshop was used to determine priority areas for improvements in documentation, performance and scalability of the workflows, accessibility of the user interface, and quality-of-life improvements to extant tools. For example, the 1.1 release of G-OnRamp includes requested improvements to Galaxy’s support for Augustus, a tool that performs comparative gene prediction [27], enabling users to limit the genomic range to search or to add extrinsic ‘hints’ for improved search specificity. Beyond this, the 1.1 release features the latest (as of this writing) versions of Galaxy (19.05), Apollo (2.4.1) and JBrowse (1.16.6). A more complete list of features is provided in Table 1B.

**Table 1B.** Feature: G-OnRamp provides….

| Processing / Analysis: |
| --- |
| The UCSC Hub Archive Creator, a tool to create genome browser archives for display with the UCSC browser |
| The JBrowse Archive Creator, a tool to create JBrowse genome browsers with Galaxy |
| An RNA-seq analysis subworkflow to process and visualize RNA-seq data |
| A BLAT alignment subworkflow to align transcript sequences from an informant genome to the target genome |
| Tools to identify repeats using WindowMasker within Galaxy |
| Input / Data Acceptance: |
| Default workflows that accept genome assemblies in fasta format, RNA-seq data in fastqsanger format, transcripts from informant genomes in GenBank or fasta formats, and proteins from informant genomes in fasta format |
| Added tools to facilitate the incorporation of results from additional gene predictors and RNA-Seq alignment tools (e.g., bigWig and BAM files) into the genome browsers produced by G-OnRamp |
| An extended Augustus tool Galaxy wrapper, exposing more functionality (e.g., ability to specify search range or add extrinsic hints)* |
| An improved Hub Archive Creator (HAC) and the JBrowse Archive Creator (JAC) tools (e.g., bug fixes, added support for new track types and custom tracks)* |
| Annotation Support: |
| Tools and a workflows to create Apollo instances from JBrowse genome browsers, and to support collaborative genome annotation using Apollo |
| Improved role-based access control in Apollo to facilitate collaborative annotation in educational settings* |
| Reporting features for instructor roles in Apollo to enable faculty to monitor student annotation progress* |
| General Ease of Use: |
| The G-OnRamp website (http://g-onramp.org), which hosts documentation, training resources and previously processed data |
| A CyVerse interaction tool to facilitate the data import and export between G-OnRamp and the CyVerse Data Store |
| JBrowse improvements to display tblastn alignments that span larger genomic regions* |
| Optimized search index strategies for feature names and descriptions in JBrowse to reduce the number of index files (e.g., Tabix-indexed GFF3 files)* |
| The ability to look up gene predictions, and the BLAST and BLAT alignments by name (e.g., RefSeq accession numbers) and by description |
| Links to external database records (e.g., at NCBI, FlyBase) for the tblastn and BLAT alignment tracks |
| Improved organization, grouping, and labeling of evidence tracks on UCSC Assembly Hubs |
| Comprehensive training materials based on feedback from the participants of the G-OnRamp beta testers workshops |
| Deployment: |
| Automated local and cloud (Amazon EC2) deployments of G-OnRamp with GalaxyKickStart — an Ansible playbook for deploying production Galaxy servers |
| A G-OnRamp image deployable via CloudLaunch (https://launch.usegalaxy.org) to enable users with limited technical expertise to run G-OnRamp on the cloud (Amazon EC2) |
| A G-OnRamp image deployable via the Amazon Web Services EC2 console |

List of major features developed for the G-OnRamp platform, and improvements made to various software components. Feature and improvement development was driven predominantly by user feedback, most of which was gathered from attendees of our biannual G-OnRamp workshops. While improvements were made throughout the cycle of G-OnRamp development, feedback from these events was a valuable aid to prioritization.
**\* Features or improvements that were developed for component services of G-OnRamp which are now generally available for those services.**

Based on the results from an anonymous survey of G-OnRamp workshop participants, we find that the overall response by users has been very good. Both researchers and educators reported that G-OnRamp has facilitated their work (Fig 5). A majority of the respondents found G-OnRamp useful in their research and/or teaching, and planned to continue to use it, including setting up new student research courses.

**Fig 5.**
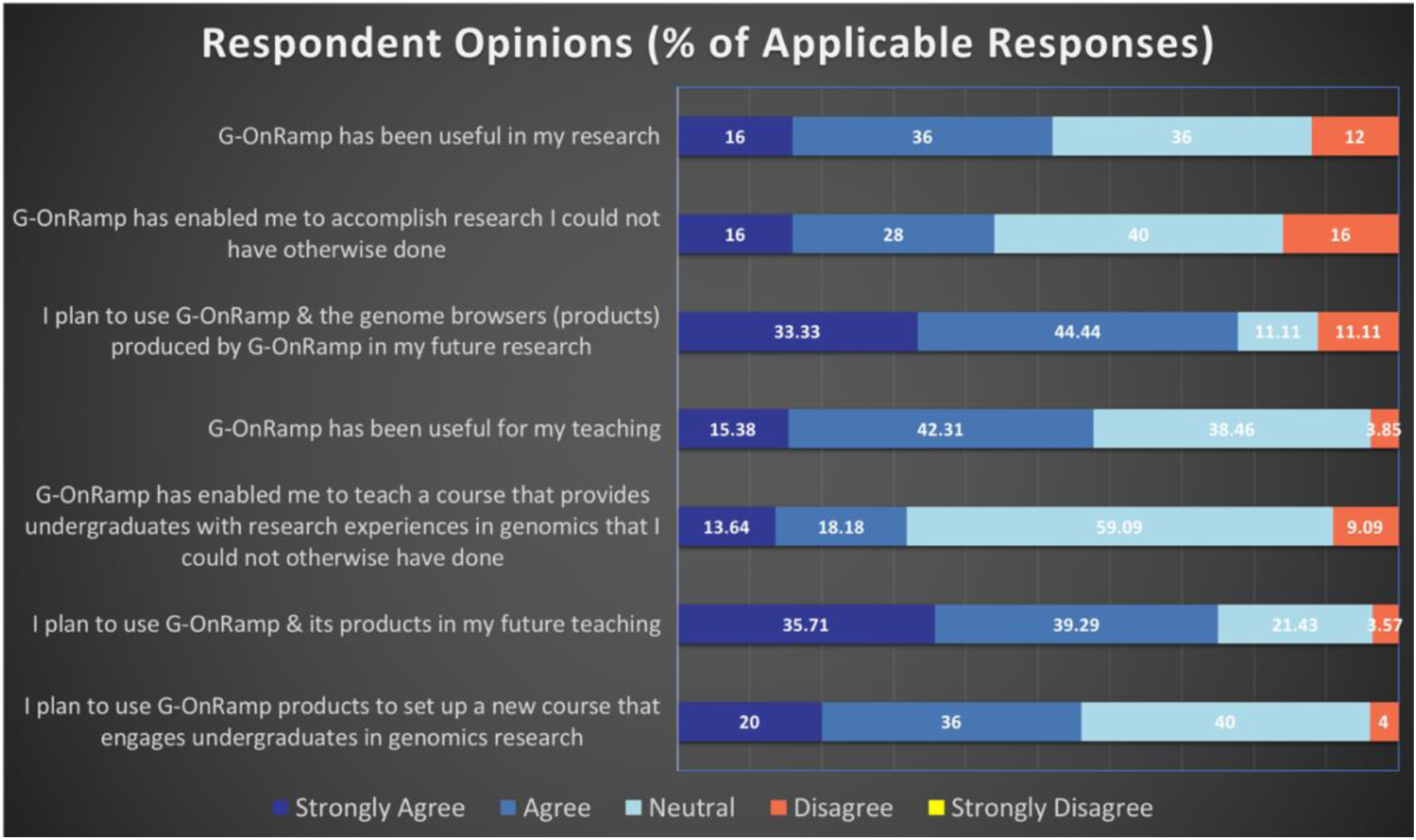
Survey responses on the utility of G-OnRamp. An anonymous survey asked respondents (N = 35 of 53 eligible) to check “strongly agree,”, “agree,” “neutral,” “disagree,” or “strongly disagree.” Participants ranged from those whose primary occupation is teaching to those managing a research support service (see Fig 4). Consequently from 20% to 38% of the participants checked “not applicable” for any given statement; these responses were removed before percentages were calculated. Overall, participants reported that G-OnRamp facilitates both research and teaching.

#### G-OnRamp in a CURE: Examining lipid synthesis pathways in parasitoid wasps

As discussed above, many bioinformatics educators have found that a genome annotation project is a good way to introduce students to genomics while providing a research experience. This can be implemented as a one-semester CURE, or as a shorter unit to provide students with an introduction to research.

Many genomics projects that can benefit from careful manual annotation will be focused on a limited set of genes. Because these genes of interest are commonly defined by a shared functional annotation or membership in a specific pathway, they are likely to be dispersed throughout the genome. In the case study presented here, the project is focused on the evolution of lipid synthesis pathways in parasitoid wasps, and so the genes of interest are defined based on their predicted functions rather than their genomic locations. This case was used to test the acceptability and utility of G-OnRamp products in the undergraduate lab.

Fig 6A illustrates the workflow underlying the creation of student annotation projects, in which the approximate locations of the genes of interest are identified in the newly sequenced genomes and assigned as student projects. Fig 6B outlines the approach taken by the student annotator, which is predicated on sequence similarity between the gene of interest in the target genome and genes from an informant genome. The difficulty of the student project primarily depends on the result of the homology search.

**Fig 6.**
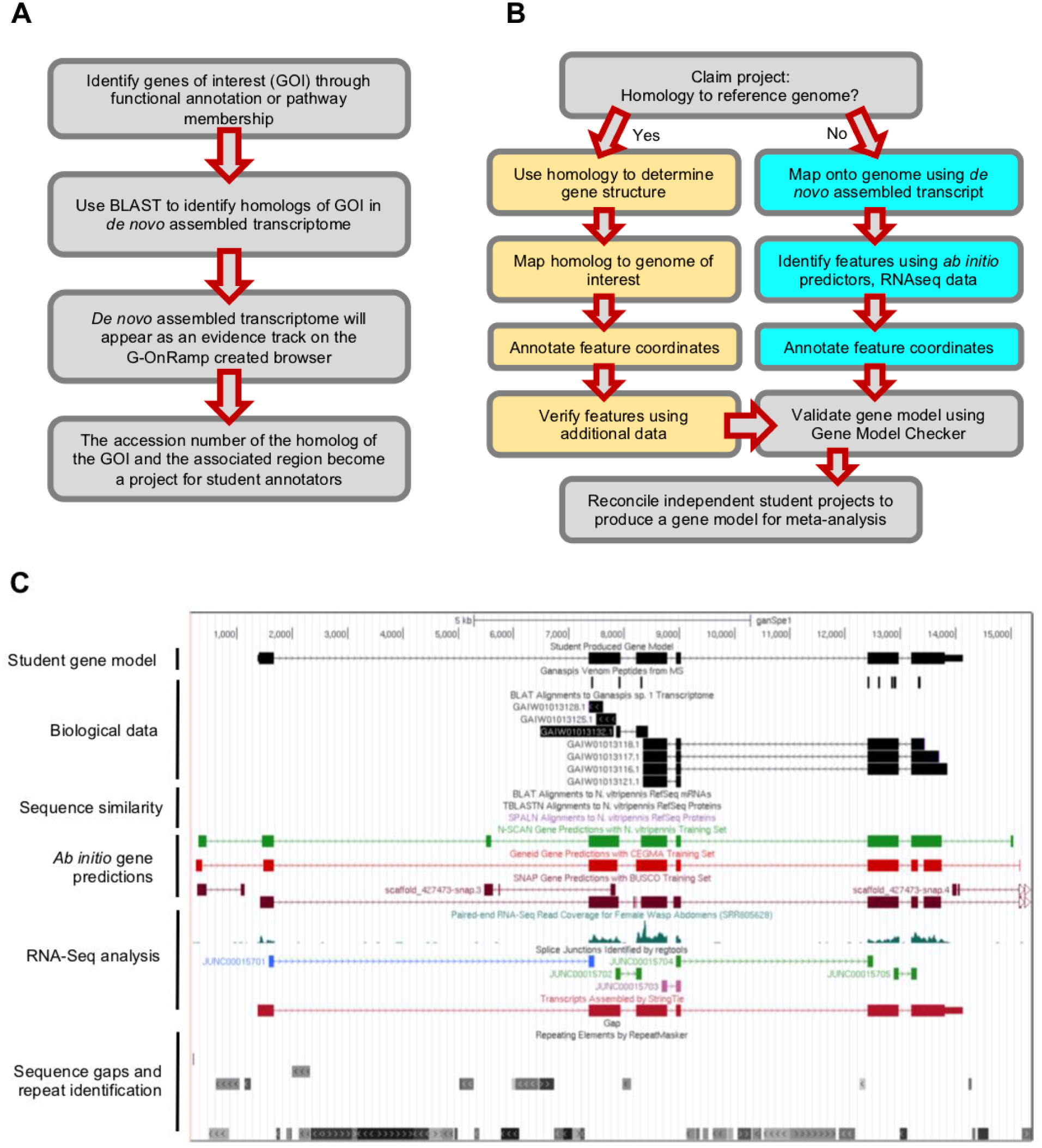
Case study: Annotation using parasitoid wasp G-OnRamp browsers. A. The workflow for identifying genes of interest and creating student annotation projects based on G-OnRamp browsers. B. The student annotation workflow. Students are assigned a project and will then work through either of the two sub-workflows depending on homology of the gene of interest to the reference genome. Boxes in yellow define the sub-workflow for genes with homology to the reference genome; cyan boxes define the sub-workflow for genes lacking homology to the reference genome. C. An example student annotation of a gene with no homology to the reference genomes (*D. melanogaster* or *N. vitripennis*). Survey respondents identified lack of homology to an informant genome as one of the main challenges in annotating new species.

A gene that aligns to an ortholog in a well-studied informant species will not be very difficult for an undergraduate to annotate, while the absence of orthologs will create a challenge. If the gene of interest has significant similarity to a gene in the informant genome, then the student annotator would construct the most parsimonious gene model compared to its putative ortholog in the informant genome. Otherwise, the student annotator would use RNA-Seq data to construct the gene model. Instructors can pre-screen projects to select those at the appropriate level of difficulty for their students.

Fig 6C illustrates an example of a student annotation of a gene that has diverged from the informant genomes (*Nasonia vitripennis* and *Drosophila melanogaster*) such that homology data are not available. The student annotator has to construct a gene model based on other lines of evidence, such as proteomics data, RNA-Seq data (*e.g.*, read coverage, *de novo* transcriptome assembly), and *ab initio* gene predictions. The flexibility of the genome browsers produced by G-OnRamp, and the annotation workflow described above, have facilitated annotation in this case, and should make comparative genomics more accessible for use in the classroom, creating opportunities to study other newly sequenced genomes.

In this pilot implementation of a CURE project using genome browsers generated by G-OnRamp, 15 faculty from the GEP designed CUREs for their students based on the parasitoid wasp research project. These faculty members came from diverse schools (Fig 7A; a full list of faculty with their schools is given in the Acknowledgements). The courses ranged from freshman/sophomore level to those that provided graduate credit. The majority were structured as a research experience.

**Fig 7.**
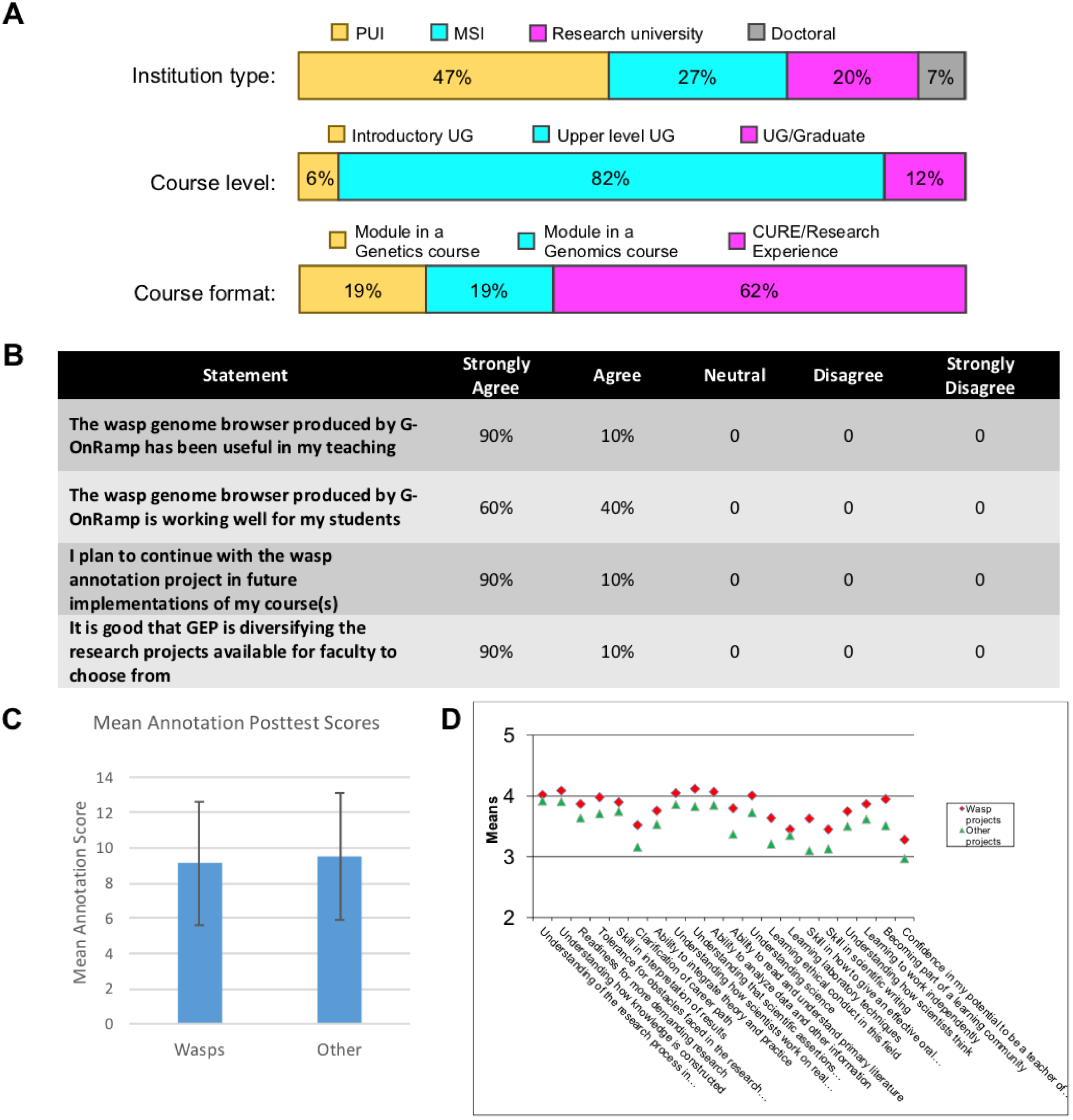
Using G-OnRamp in a CURE. Classroom implementation with G-OnRamp browsers. A. Implementations of the parasitoid wasp project during 2017-2018 and 2018-2019 characterized by institution type (n = 15), course level (n = 16) and course format (n = 16). Abbreviations: PUI = Primarily undergraduate institution, MSI = Minority-serving institutions, UG = undergraduate, CURE = Course-based Undergraduate Research Experience. B. Results from a survey of faculty who have used a G-OnRamp-generated browser in a course. Participants were asked to respond on a 5-point Likert Scale with N.A. as an option; of the 14 faculty responding to this portion of the survey, the four checking “NA” for these questions were removed before calculating percentage responses, giving n = 10. Responses are shown by percentage of respondents. C. Mean annotation post-course test scores: The mean for the Wasp group is 9.1 (N = 173; SD = 3.6) and the mean for the other GEP students is 9.5 (N = 1185; SD = 3.5). The difference is not significant (bars represent the means; error bars represent one standard deviation). D. Responses to the SURE survey questions: the means for the wasp project students are in red (N ranges from 181 to 195, as some students did not answer all questions) and the means for the other GEP students (working in Drosophila) are in green (N ranges from 1200 to 1270). For some items the wasp group scores significantly higher than the comparison group; however, these results should be interpreted with caution, given the small sample size.

Responses from an anonymous survey show that most faculty found that the wasp genome browser produced by G-OnRamp worked well for their students, and was generally useful in teaching (Fig 7B). Faculty members who responded to the survey all planned to continue involving their students in the parasitoid wasp project the following year, and all applauded the effort by the GEP/Galaxy partnership to support genomics research broadly.

Direct assessment of the students engaged in a parasitoid wasp CURE was obtained by comparing the responses of this group to those of GEP students as a whole, looking at pooled data from 2017–2018 and 2018–2019. The results show no significant difference in student attainment as exhibited by post-course quiz scores (Fig 7C), indicating that the G-OnRamp-produced genome browsers and the wasp research project are as effective as the UCSC mirror *Drosophila* genome browsers and Muller F element research project in teaching the fundamentals of eukaryotic genes and genomes. Interestingly, there is a small increase in the responses to the SURE survey questions [28], which ask students to self-report perceived gains in the understanding of how science is done and their acquisition of research skills (Fig 7D). This suggests that G-OnRamp can increase student and faculty enthusiasm for genomics research by enabling a variety of projects. Eventually we hope to see multiple collaborative annotation projects that would allow all faculty to participate in a project according to their research interests.

#### Using G-OnRamp on your own

Steps for acquiring and deploying G-OnRamp, like the platform itself, minimize technical complexity and accelerate data analysis activities. The two principal methods of deployment meet different user needs: 1.) a VirtualBox virtual appliance for small-scale local testing and training and 2.) an Amazon Machine Image (AMI) for cloud-based production deployments. Users can launch the G-OnRamp AMI on Amazon Web Services (AWS) via the CloudLaunch web application (https://launch.usegalaxy.org/; Table 2).

**Table 2.**
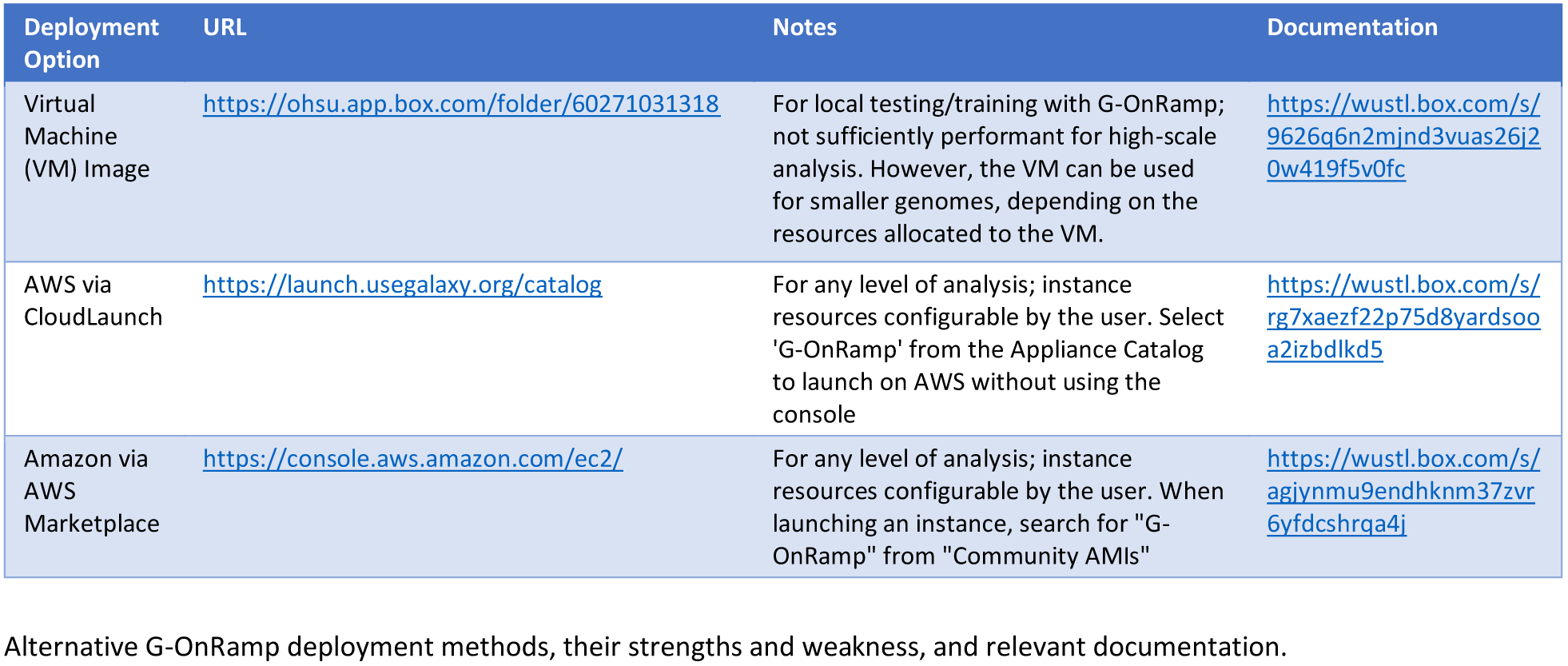
Deployment options.

For more fine-grained control of the installation and launch of G-OnRamp, the scripts used to create the two principal deployment options are open-source and available on GitHub (https://github.com/goeckslab/gonrampkickstart). This option provides much greater control, but comes with additional complexity that requires technical expertise. For more complex deployment configurations within the AWS infrastructure, a G-OnRamp image can be found under “Community AMIs” when launching an Elastic Cloud Compute (EC2) instance.

## Conclusion

The importance and efficacy of providing undergraduates with a research experience is widely accepted. While it is difficult to identify the impact of research *per se* [29], students engaged in a CURE are reported to be both retained in the sciences and to graduate within six years at a higher frequency than matched students who do not have this experience [30]. CUREs in bioinformatics have many advantages, both practical and pedagogical: infrastructure costs are low (only requires computers and Internet connectivity), and there is a large and growing pool of publicly available data, along with tools to manage and analyze that data (*e.g.*, Galaxy, CyVerse). Because no physical lab is required, access is 24/7, and there are no lab safety issues; this situation lends itself to peer instruction, an important multiplier. Perhaps most important, student mistakes are inexpensive in time and money, as the annotation process can be quickly reiterated, problems explored, and investigations taken to the next level. Recognizing these advantages, a growing number of faculty groups have emerged over the last decade to organize CUREs that include collaborative genome annotation [8, 31, 32, 33]. Recently, several of these groups have come together to form a Genomics Education Alliance (<u>GEA;</u> https://qubeshub.org/community/groups/gea/), which seeks to support this effort by creating a common, well-maintained platform with common curriculum and tools [34]. G-OnRamp removes one bottleneck to CURE growth in bioinformatics by facilitating creation of the genome browsers needed for collaborative genome annotation projects. The G-OnRamp survey results and the parasitoid wasp pilot project have shown G-OnRamp to be a useful tool for researchers and educators alike.

## Acknowledgements

We thank the GEP faculty members and their students who participated in the parasitoid wasp project during the last two years: Cindy Arrigo (New Jersey City University), Rebecca Burgess (Stevenson University), Thomas Giarla (Siena College), Rivka Glaser (Stevenson University), Shubha Govind (City College, CUNY), Adam Haberman (University of San Diego), Christopher Jones (Moravian College), Lisa Kadlec (Wilkes University), Adam Kleinschmit (University of Dubuque), Leocadia Paliulis (Bucknell University), Srebrenka Robic (Agnes Scott College), Michael Rubin (University of Puerto Rico at Cayey), Sheryl Smith (Arcadia University), Joyce Stamm (University of Evansville), and Melanie Van Stry (Lane College). Development of G-OnRamp was supported by NIH grant 1R25 GM119157 “A genome browser on-ramp to engage biologists with big data” awarded to SCRE; the work on parasitoid wasps is supported by NIH grants 1R35 GM133760 and 1R03 AG063314 to NTM.

## Notes

http://g-onramp.org/

